# Tonic feedback motor commands predict visuomotor learning

**DOI:** 10.64898/2026.01.30.702978

**Authors:** Yuto Makino, Toshiki Kobayashi, Daichi Nozaki

**Affiliations:** Center for Information and Neural Networks, National Institute of Information and Communications Technology, Osaka 565-0871, Japan; Division of Physical and Health Education, Graduate School of Education, The University of Tokyo, Tokyo 113-0033, Japan; Japan Society for the Promotion of Science, Tokyo 102-0083, Japan; Sony Computer Science Laboratories, Inc., Tokyo 141-0022, Japan; Integrated Open Systems Unit, Okinawa Institute of Science and Technology Graduate University, Okinawa 904-0495, Japan; UTokyo Sports Science Initiative, The University of Tokyo, Tokyo 153-8902, Japan

## Abstract

When a movement error occurs, the motor system updates its commands to improve performance on subsequent trials. A prominent feedback error learning hypothesis proposes that the feedback response that corrects movement within a trial serves as a teaching signal for the learning response, observed as changes in motor commands on the next trial. However, how the temporal pattern of the feedback response influences the learning response remains unclear. Here, we introduce an experimental paradigm that directly compares the temporal patterns of feedback and learning responses across different patterns of visual error. We show that although the feedback response closely tracks the temporal pattern of the visual error, this temporal pattern is not transferred to the learning response. Instead, the amplitude of the feedback response during the holding period, which reflects the temporal history of the visual error, strongly predicts the magnitude of the learning response. These findings suggest that information reflected in the tonic component of feedback responses is closely linked to the scaling of subsequent motor-command updates.

## Introduction

Accurate and consistent movements rely on the motor system’s ability to correct movement errors. Motor adaptation, the process by which motor commands are updated based on sensory error information, has been extensively studied using perturbations during arm reaching tasks. In these paradigms, lateral deviations of the hand or cursor trajectory are typically treated as scalar measures of both motor commands and movement errors. This framework has enabled systematic investigation of key properties of motor adaptation, including the time course of adaptation and deadaptation [1], interference between opposing perturbations [2], generalization across targets [3–4], context dependency [5], and the presence of multiple timescales of motor memory [6].

Despite these advances, two fundamental questions remain. First, sensory error information is represented in sensory space rather than motor command space, and therefore cannot be directly used to update motor commands [7–9]. Second, treating motor commands as scalar values fails to capture their temporal dynamics. The feedback error learning hypothesis [9] has been proposed as a mechanism that addresses these issues. According to this hypothesis, the motor commands that correct a movement within a trial, referred to as feedback responses (FBRs), serve as teaching signals for updating motor commands expressed in subsequent trials, referred to as learning responses (LRs).

Albert and Shadmehr [10] provided behavioral evidence supporting this idea by comparing muscle activity during feedback responses in force field perturbation trials with learning responses in subsequent trials. They reported that learning responses resembled feedback responses temporally advanced by approximately 125 ms, and that larger feedback responses led to larger learning responses, suggesting that feedback responses may act as teaching signals. However, other studies have shown that feedback responses follow the temporal characteristics of the perturbation, whereas such temporal features are not preserved in learning responses [11–12]. Moreover, our recent work [13] demonstrated a dissociation between feedback and learning responses when multiple or variable visual errors were imposed. Together, these findings challenge the notion that feedback responses universally serve as direct teaching signals.

Thus, a systematic reinvestigation of the relationship between feedback and learning responses is essential for understanding how feedback responses contribute to motor learning. To address this issue, we combined a single trial adaptation paradigm [3,10,13–15] with a force channel method originally developed to quantify feedback responses to visual errors [16–17]. This approach enables direct comparison of the temporal profiles and magnitudes of feedback and learning responses elicited by different visual error patterns, while minimizing changes in movement kinematics.

The present study aimed to characterize the relationship between feedback responses and subsequent learning responses, focusing on their temporal profiles and amplitudes. We addressed three questions. First, how does the learning response depend on the onset timing of the feedback response? Second, how does the amplitude of the feedback response influence the amplitude of the learning response? Third, how is the learning response shaped when the temporal pattern of the feedback response is drastically altered? Together, these questions allowed us to characterize the temporal and quantitative relationship between feedback responses and subsequent learning responses across different visual-error patterns.

## Results

### Experimental paradigm to compare the FBR and LR

A total of 33 healthy participants took part in the study (Experiments 1–3). They were instructed to move a visual cursor toward a frontal target by moving the handle of a robotic manipulandum (KINARM). Each participant repeated a cycle consisting of three consecutive force channel trials (baseline, perturbation, and probe trials) during which the handle trajectory was constrained along the straight path [18], followed by two ordinary (i.e., without force channel) reaching trials to washout the possible learning effect (Fig. 1a). In the baseline trial (Fig. 1b), the lateral force exerted against the force channel was measured to determine a baseline force when no perturbation was imposed on the cursor. In the subsequent perturbation trial (Fig. 1c), the cursor position was shifted laterally either to the left or right (i.e., cursor shift perturbation; [16–17]). The FBR was obtained as the lateral force induced by this cursor shift, subtracted by the baseline force. In the following probe trial (Fig. 1d), the LR elicited by the preceding perturbation was obtained as the lateral force, again subtracted by the baseline force. During this probe trial, the cursor was invisible to prevent any online movement correction. For analysis, the temporal profiles of FBR and LR were collapsed across rightward and leftward cursor shift perturbations (Fig. 1e).

**Figure 1:**
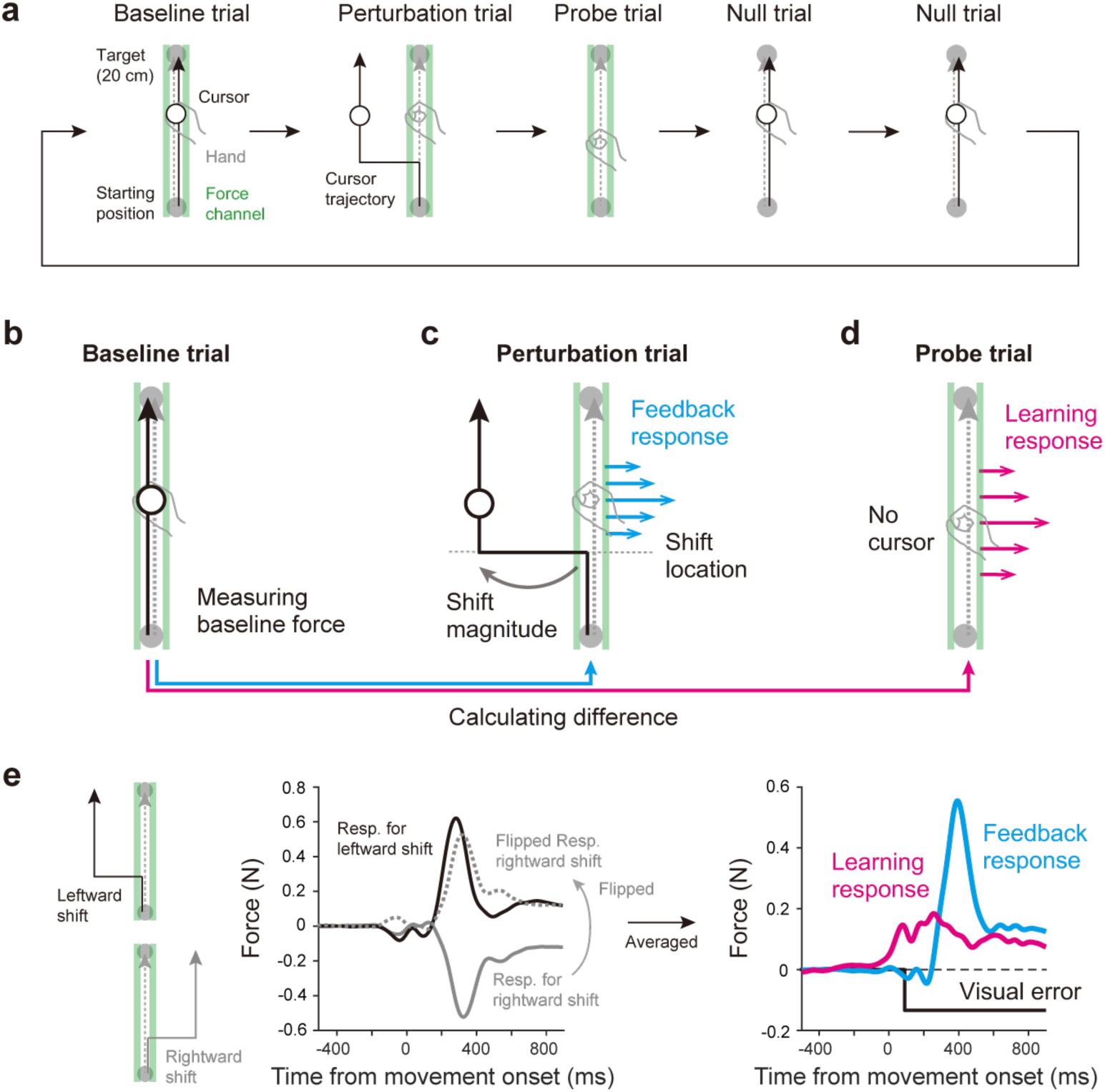
Procedures of the experimental paradigm. We employed a cursor shift perturbation paradigm to reinvestigate the relationship between feedback responses (FBRs) and learning responses (LRs). **A,** Participants repeatedly performed a block consisting of one baseline trial, one perturbation trial, one probe trial followed by two null trials. **B,** Baseline trials. Forces exerted against the force channel were measured to quantify baseline motor output. **C,** In perturbation trials, the cursor was shifted laterally by manipulating both location and magnitude of the shift. This manipulation allowed the imposition of arbitrary temporal patterns of visual error. Hand trajectories were constrained using the force channel. **D,** Probe trials were used to measure the LR elicited for the visual error imposed in the preceding perturbation trial. Temporal patterns of FBRs and LRs were baseline-subtracted using forces measured during baseline trials. **E,** Temporal profiles of responses to rightward and leftward cursor shift perturbations were collapsed (left) to enable direct comparison of the motor commands underlying FBRs and LRs for a given temporal pattern of visual error.

This experimental paradigm offers several advantages over conventional methods for investigating the relationship between visual errors, FBRs, and LRs. First, the cursor shift perturbation allowed us to impose arbitrary temporal patterns of visual errors by manipulating the timing and magnitude of the cursor shift (Fig. 1c). Second, the temporal profiles of motor commands associated with both FBR and LR could be directly compared, as they were derived from forces measured against the force channel and were therefore not confounded by changes in movement kinematics.

### Experiment 1: Effect of visual perturbation onset on the temporal profiles of FBR and LR

The feedback error learning hypothesis proposes that FBRs serve as teaching signals for LRs. Consistent with this idea, a previous study reported that LRs are time shifted versions of FBRs, with a temporal advance of approximately 125 ms [10].

Experiment 1 (10 participants; seven men and three women) examined how the onset timing of visual perturbations influences the relationship between the temporal profiles of FBRs and LRs. In the perturbation trial, the cursor was shifted laterally by 3 cm either to the right or left at different distances from the starting position (Fig. 2a; 1, 6, 11, and 16 cm). In addition, one condition involved no cursor shift, resulting in a total of nine perturbation conditions (two shift directions × four shift locations + one no shift condition).

**Figure 2:**
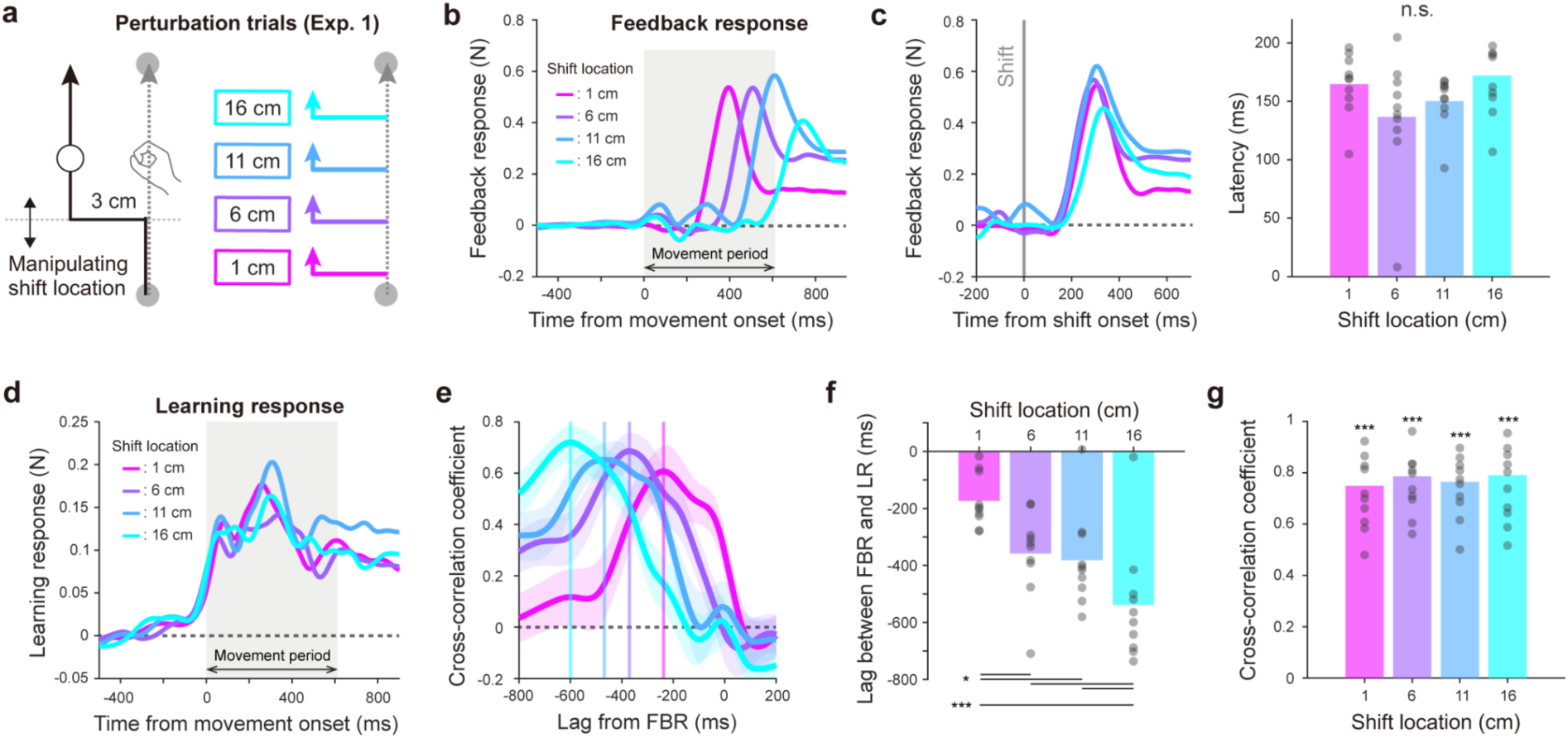
Relationship between feedback and learning responses in Experiment 1. **a**, In the perturbation trial, the cursor shift location was manipulated from 1 cm to 16 cm from the starting position, while the cursor shift magnitude was fixed at 3 cm. **b,** Temporal profiles of the feedback responses (FBRs) in Experiment 1, aligned to movement onset. **c,** FBRs emerged with a latency of approximately 160 ms relative to movement onset. Latencies were estimated using receiver operating characteristic (ROC) analysis. **d,** Temporal profiles of learning responses (LRs) in Experiment 1. **e-g,** Cross-correlation analysis was used to evaluate temporal similarity and relative timing between FBRs and LRs (**e**). The temporal shift between the FBRs and LRs was defined as the time lag that maximized the individual cross-correlation (**f**), and temporal similarity was quantified as the cross-correlation coefficient at this optimal time lag (**g**). Shaded areas represent the standard error of the mean across participants. Asterisks indicate statistically significant effects (* p < 0.05, ** p < 0.01, *** p < 0.001).

Figure 2b shows the FBRs, obtained by subtracting the lateral forces measured during the baseline trial from those measured during the perturbation trial. Although participants were instructed not to use any explicit strategy, the cursor shift elicited implicit FBRs in the direction opposite to the cursor shift, consistent with compensatory corrections. The latency of FBRs was estimated using a receiver operator characteristic analysis [19–20]. This analysis revealed that feedback responses emerged 155.9 ± 13.9 ms after the cursor shift onset, with no significant differences across shift locations (Fig. 2c; F(3,27) = 1.688, p = 0.193, one-way repeated-measures ANOVA). This latency is consistent with previous reports of rapid visually driven feedback responses [17,21–23].

If LRs were fixed time shifted copies of FBRs [10], the onset of LRs should vary systematically with the onset of FBRs, and their temporal profiles should closely match. However, the lateral forces measured during the probe trial revealed a different pattern. Figure 2d illustrates the LRs, computed as the difference in lateral force between the probe and baseline trials, showing that LRs consistently emerged just before movement onset, regardless of the cursor shift location.

To quantify the temporal relationship between FBRs and LRs, we computed the cross-correlation between the two responses (Fig. 2e). One-way repeated-measures ANOVA revealed that the temporal shift corresponding to the maximum cross-correlation varied significantly with the perturbation onset location (Fig. 2f; F(1.682, 15.135) = 11.341, p = 0.001, Greenhouse Geisser correction). In contrast, Fisher transformed cross-correlation coefficients were significantly greater than zero for all perturbation conditions at the lag that maximized the correlation (Fig. 2g; 1 cm: t(9) = 9.182, p < 0.001; 6 cm: t(9) = 8.956, p < 0.001; 11 cm: t(9) = 11.747, p < 0.001; 16 cm: t(9) = 8.380, p < 0.001), indicating similar temporal profiles across responses. Together, these results indicate that although FBRs and LRs share similar temporal profiles, the relative timing between them is not fixed. Instead, LRs consistently emerge just before movement onset, independent of the onset timing of the FBRs.

### Experiment 2: Effect of the size of visual error on the quantitative relationship between FBR and LR

In Experiment 1, the magnitude of the cursor shift was fixed at 3 cm while its onset location was varied. Experiment 2 (12 participants; six men and six women) examined how the magnitude of the cursor shift, while keeping its location constant, influenced FBRs and LRs. In the perturbation trials, the cursor was shifted laterally by 0.4, 0.8, 1.2, 1.6, 2.0, or 3.0 cm, all applied at a location 1 cm from the starting position (Fig. 3a).

**Figure 3:**
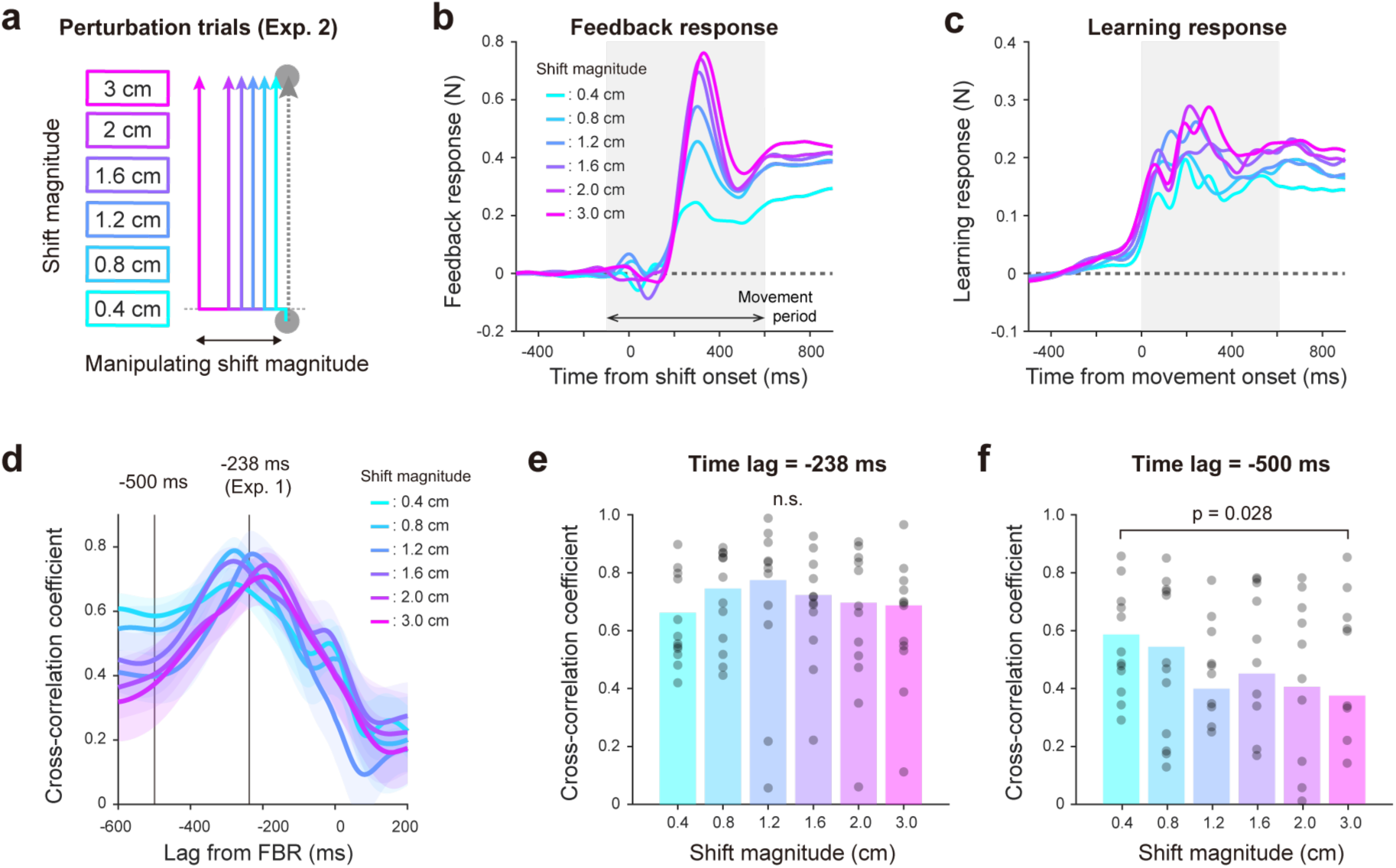
Temporal patterns of feedback and learning responses across different cursor shift magnitudes in Experiment 2. **a**, In perturbation trials, the shift magnitude was varied from 0.4 cm to 3 cm. **b,** Temporal patterns of the feedback responses (FBRs) in Experiment 2. **c,** Temporal profiles of learning responses (LRs) in Experiment 2. **d,** Cross-correlation analysis between FBRs and LRs used to evaluate their temporal similarity. The plot shows the cross-correlation coefficient as a function of time lag. **e,** Cross-correlation coefficient at the time lag that maximized the average cross-correlation function in Experiment 1 (−238 ms). **f,** Cross-correlation coefficient at a fixed time lag of -500 ms. A significant modulation of correlation coefficients was observed across error magnitudes. Shaded areas represent the standard error of the mean across participants.

Figure 3b illustrates the temporal patterns of FBRs across different cursor shift magnitudes. As observed in Figure 2, the FBRs consisted of an early phasic component followed by a later tonic component. Franklin et al. [23] reported that the magnitude of the FBR immediately after cursor shift onset is linearly modulated by the magnitude of the cursor shift. Consistent with this finding, the amplitude of the early phasic component increased systematically with increasing cursor shift magnitude. In contrast, the amplitude of the later tonic component, observed near movement offset, appeared to saturate when the cursor shift magnitude exceeded 0.8 cm. Figure 3c illustrates the temporal patterns of LRs across different cursor shift magnitudes. In contrast to FBRs, the waveforms of LRs remained largely similar across conditions, while their amplitudes showed only modest modulation with cursor shift magnitude.

To further characterize how the temporal patterns of FBRs and LRs vary with error magnitude, we performed cross-correlation analyses for each cursor shift magnitude (Fig. 3d). This analysis revealed that the overall shape of the cross-correlation function changed systematically with error magnitude. To quantify these differences, we evaluated cross-correlation coefficients at two specific time lags between the FBRs and LRs: a fixed lag corresponding to the maximum cross-correlation observed in the 1 cm condition of Experiment 1 (Fig. 3e; -238 ms), and a longer lag of -500 ms (Fig. 3f). A two-way repeated-measures ANOVA revealed a significant interaction between time lag and error magnitude (F(5,55) = 3.061, p = 0.017). We then performed simple main effects analyses at each time lag. These analyses showed that the main effect of error magnitude was significant at the longer lag (Fig. 3f; -500 ms: F(5,55) = 2.734, p = 0.028), but not at the shorter lag (Fig. 3e; -238 ms: F(5,55) = 0.891, p = 0.494). This selective modulation of temporal similarity at longer lags indicates that the temporal profile of the FBR changes with increasing error magnitude, potentially reflecting the recruitment of distinct feedback-related components. In contrast, the temporal patterns of LRs remained relatively invariant across error magnitudes.

To quantitatively evaluate how response components vary with error magnitude, we decomposed the waveforms of the FBRs and LRs, [R(t)], into phasic and tonic components using the following equation (Figs. 4a, c):

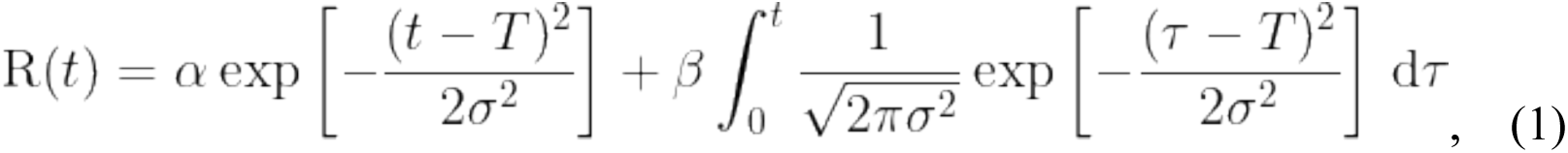

where α, β, σ and *T* are constants. Note that α and β represent the amplitudes of early phasic and late tonic components, respectively. This formulation was inspired by previous studies suggesting that motor commands during reaching consist of two components: a movement related signal and a holding related signal, with the latter generated by integrating the former over the course of the movement [24–25].

**Figure 4:**
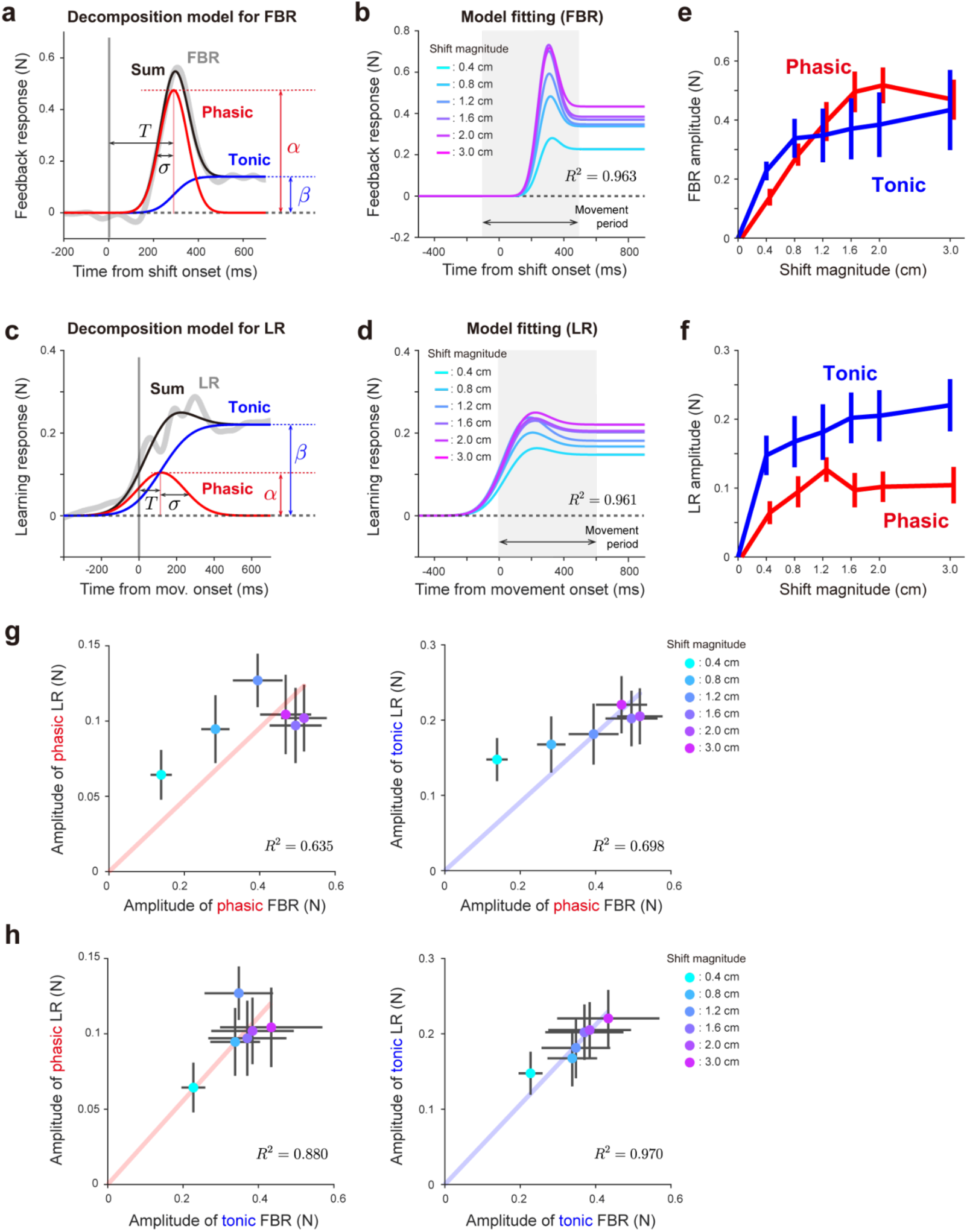
Decomposition of feedback and learning response temporal profiles. **a**, Temporal profiles of feedback responses (FBRs) were decomposed into two components: an early phasic component and a late tonic component. **b,** Decomposition model reproduced the temporal profiles of FBRs in Experiment 2. **c-d,** The same decomposition procedure was applied to the temporal profiles of the learning responses (LRs). **e,** Modulation of the amplitudes of the phasic and tonic components of the FBR as a function of error magnitude. **f,** Modulation of the amplitudes of the phasic and tonic components of the LR as a function of error magnitude. Error bars represent the standard error of the mean across the participants. **g-h,** Linear regression analyses examining the quantitative relationship between FBRs and LRs using the model *y = ax*, where *x* represents the amplitude of the phasic (**g**) and tonic components (**h**) of the FBR, and *y* represents the amplitude of the corresponding phasic (left) and tonic components (right) of the LR. Error bars represent the standard error of the mean across the participants.

We first fitted the FBRs, aligned to the onset of the cursor shift and averaged across participants, using Eq. 1. The fitted model accurately reproduced the experimentally observed temporal patterns of the FBR (Fig. 4b, σ = 0.9627, σ *=* 59, *T* =289). Assuming that the temporal parameters of the model (σ and *T*) were approximately consistent across participants, we then refitted the feedback responses for each individual while estimating only the amplitudes of the phasic and tonic components. Notably, allowing the temporal parameters to vary freely did not substantially affect the results reported below (Supplementary Figure. 1).

We applied the same analysis to the LRs aligned to movement onset, which yielded similarly good fits (Fig. 4d; *R*^2^ = 0.961, σ = 126, *T* = 118). Figures 4e and 4f show how the amplitudes of the phasic and tonic components of both responses varied with error magnitude. Consistent with the temporal patterns shown in Figs. 3b and 3c, the phasic component of the FBR increased approximately linearly with error magnitude up to 1.8 cm. In contrast, the tonic component of the FBR and both the phasic and tonic components of the LR saturated at approximately 0.8 cm (Figs. 4e, f).

Based on the near linear scaling of the phasic FBR component and the early saturation of the other components, we hypothesized that LR amplitudes would be more closely associated with the tonic component of the FBR than with the phasic component. To test this hypothesis, we performed linear regression analyses using the model *y = ax*, where *x* represents the amplitude of each FBR component and *y* represents the corresponding component of the LR. As shown in Figure 4g, the phasic FBR component exhibited only moderate predictive power (R² = 0.635 for the phasic LR component and R² = 0.698 for the tonic LR component). In contrast, when the tonic FBR component was used as the predictor (Fig. 4h), both relationships were markedly stronger (R² = 0.880 for the phasic LR and R² = 0.970 for the tonic LR), indicating a higher degree of linearity. Together, these results indicate that the tonic component of the FBR provides the strongest predictor of LR modulation across different error magnitudes.

### Experiment 3: Effect of biphasic temporal patterns of visual error on the FBR and LR

Although the phasic and tonic components of FBRs and LRs were differently modulated by error magnitude in Experiment 2, the overall temporal profiles of both responses remained largely similar across Experiments 1 and 2, regardless of cursor shift position or error magnitude. This similarity may reflect the use of stepwise cursor shift perturbations in both experiments, which limited the range of temporal patterns of visual error and may have constrained dissociation between the two responses.

To more directly test whether the temporal patterns of the FBR and LR could be dissociated, we introduced more complex temporal patterns of visual error in Experiment 3 (11 participants: six men and five women). Specifically, three different temporal patterns of cursor shift perturbation were employed (Fig. 5a). In all perturbation conditions, the cursor was first shifted laterally by ±3 cm at the 1 cm location. Following this first shift, the cursor position was either maintained (maintained condition), returned to its original position (removed condition), or shifted to the opposite side (reversed condition) (Fig. 5a). The second cursor shift, when present, occurred at one of three locations (6, 11, and 16 cm).

**Figure 5:**
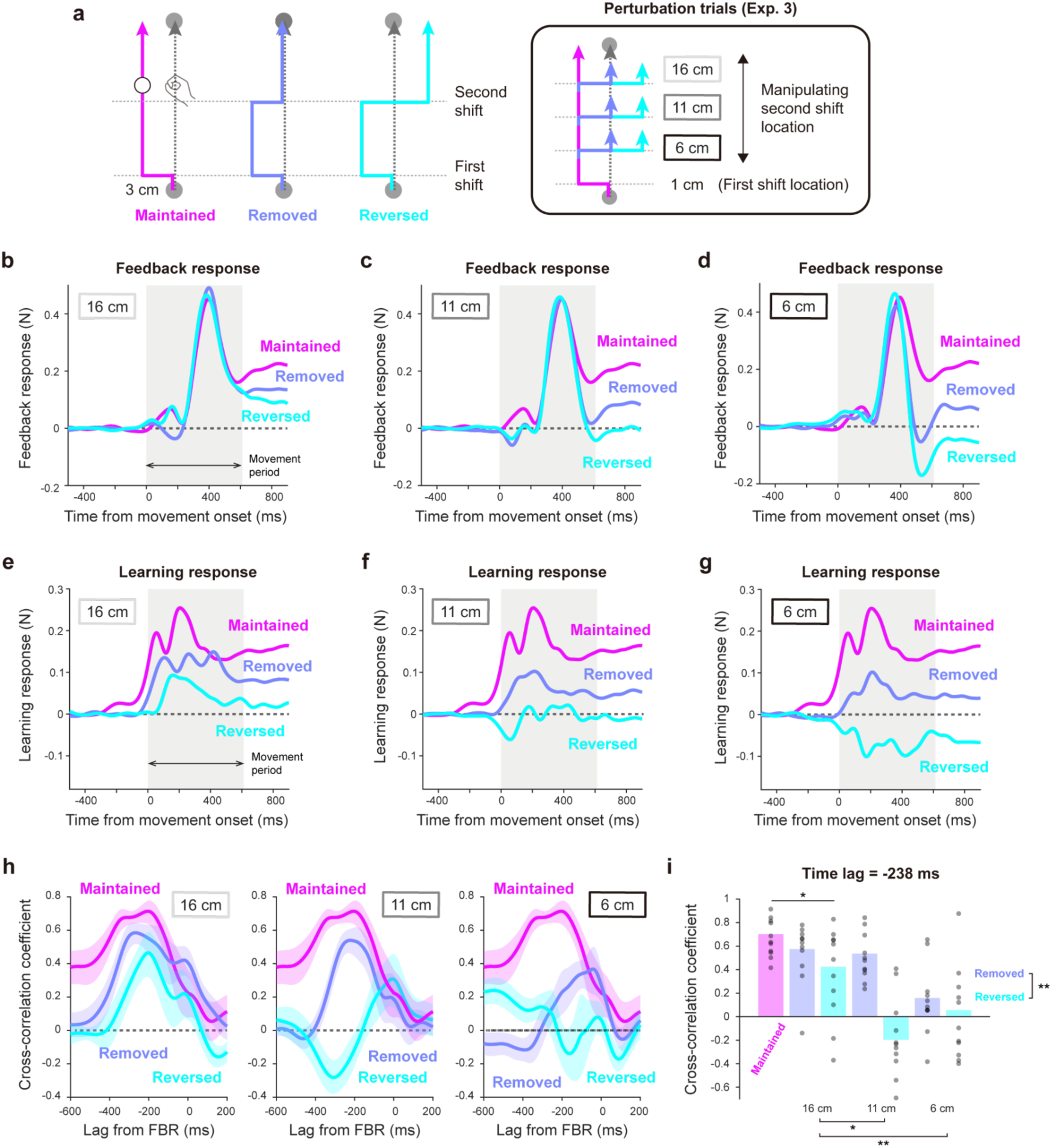
Relationship between the temporal patterns of feedback and learning responses in Experiment 3. **a**, In perturbation trials, the cursor was shifted laterally twice during a single movement. Three temporal patterns of visual error were tested, and the location of the second shift was varied from 6 cm to 16 cm from the starting position. **b-d,** Temporal profiles of the feedback responses (FBRs) when the second shift occurred at 16 cm (**b**), 11 cm (**c**), and 6 cm (**d**). **e-g,** Temporal patterns of the learning responses (LRs) when the second shift occurred at 16 cm (**e**), 11 cm (**f**), and 6 cm (**g**). **h,** Cross-coefficient function showing the relationship between the cross-correlation coefficient and time lag between FBRs and LRs when the second shift occurred at 16 cm (left), 11 cm (middle), and 6 cm (right). **i,** Cross-correlation coefficient evaluated at the time lag that maximized the cross-correlation function in Experiment 1 (−238 ms). Shaded areas represent the standard error of the mean across participants. Asterisks indicate statistically significant effects (* p < 0.05, ** p < 0.01, *** p < 0.001).

### The temporal pattern of the LRs differs from that of the FBRs

Figures 5b–d illustrate the temporal profiles of the FBRs to cursor shift perturbations in Experiment 3. FBRs were sensitive not only to the initial cursor shift but also to the second shift. For example, when the second cursor shift occurred late in the movement at 16 cm, the feedback responses during the movement period were nearly identical across conditions, with differences emerging primarily during the holding period (Fig. 5b). As the second shift was applied earlier in the movement, these differences became increasingly pronounced. Notably, when the second shift occurred at 6 cm (Fig. 5d), the FBR returned to baseline in the removed condition and became negative in the reversed condition.

In contrast to the FBRs, the LRs did not follow the temporal patterns of the cursor shift perturbations (Figs. 5e–g) and therefore did not resemble the FBRs. For example, when the second cursor shift occurred at 6 cm in the reversed condition, the biphasic pattern observed in the FBR (Fig. 5d) differed markedly from the temporal profile of the LR (Fig. 5g). To quantify this dissociation, we computed cross-correlation functions between the FBR and LR (Fig. 5h) and assessed similarity using the cross-correlation coefficient at a fixed time lag (Fig. 5i). This lag corresponded to the time yielding maximum cross-correlation in the 1 cm condition of Experiment 1 and was used as a reference, as in Experiment 2.

As in Experiment 1, the cross-correlation coefficient in the maintained condition was approximately 0.7, indicating moderate similarity between FBRs and LRs. However, this similarity was significantly modulated by both the temporal structure of the second shift and its location (Fig. 5i). Focusing on conditions where the second shift occurred at 16 cm, a one-way repeated-measures ANOVA revealed a significant main effect of perturbation condition on the cross-correlation coefficient (F(2,20) = 3.633, p = 0.045). Post hoc testing indicated that similarity was significantly lower in the reversed condition than in the maintained condition (t(10) = 2.695, p = 0.042, Holm-corrected). These results indicate that even when the second shift occurred late in the movement, altering the temporal structure of the visual error reduced the similarity between FBR and LR.

Focusing further on the removed and reversed conditions, a two-way repeated-measures ANOVA revealed a significant interaction between error pattern and second shift location (F(2,20) = 4.569, p = 0.023). Post hoc tests showed significant main effects of both error pattern (t(10) = 4.122, p = 0.002, Holm-corrected) and shift location (11 cm vs. 16 cm: t(10) = 2.784, p = 0.023; 6 cm vs. 16 cm: t(10) = 3.493, p = 0.007, Holm-corrected). Together, these results provide strong evidence that the temporal pattern of the LR does not consistently resemble that of the FBR.

### The tonic component of the FBR predicts LR modulation induced by the second shift

Focusing on conditions in which the second cursor shift was imposed at 16 cm, the FBRs were nearly identical during the movement period and diverged only slightly during the holding period (Fig. 5b). Despite this subtle difference, the amplitude of the LR in the subsequent trial differed markedly across conditions (Fig. 5e), consistent with the results of Experiment 2 and suggesting a close relationship between the tonic component of the FBR and LR amplitude. To quantify the coupling between FBRs and LRs attributable specifically to the second shift, we isolated its contribution by subtracting the FBRs and LRs in the maintained condition (no second shift) from those in the removed and reversed conditions (Figs. 6a–b, inset). We then fitted the decomposition model to these differential waveforms of the FBR and LR to test our hypothesis.

**Figure 6:**
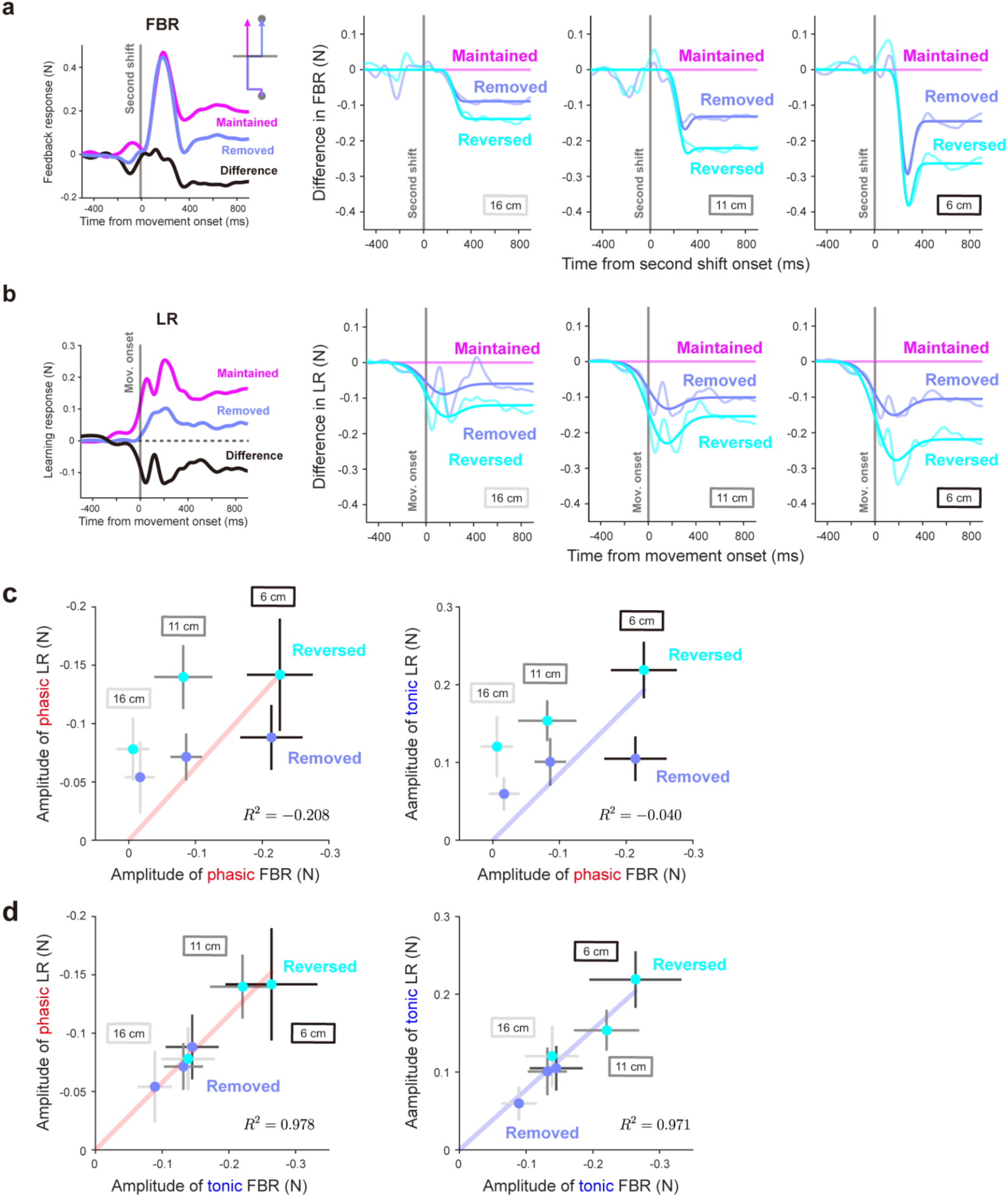
Temporal relationship between feedback and learning responses in Experiment 3. **a-b**, To isolate the effect of the second cursor shift on feedback (FBRs) and learning responses (LRs), differential force profiles were computed by subtracting the responses in the maintained condition from those in the removed and reversed conditions (insets). The decomposition model successfully reproduced the differential force profiles of the feedback response (**a**) and learning response (**b**) for the second shift when it occurred at 16 cm (left), 11 cm (middle), and 6 cm (right). Solid lines indicate model predictions, and dashed lines indicate experimental data. **c,** Linear regression analysis examining the quantitative relationship between the phasic component of the FBR and the amplitudes of the phasic (left) and tonic (right) components of the LR for the second cursor shift, following the same procedure as in Experiment 2. **d,** Linear regression analysis examining the relationship between the tonic component of the FBR and the amplitudes of the phasic (left) and tonic (right) components of the LR for the second cursor shift. Error bars represent the standard error of the mean across participants.

The decomposition model successfully reproduced the differential force profiles of both FBRs (Fig. 6a; *R*^2^ = 0.913, σ = 47, T = 263) and LRs (Fig. 6b; *R*^2^ = 0.815, σ = 131, T = 97). We next examined how the second cursor shift affected the phasic and tonic components of FBRs and LRs. Following the approach used in Experiment 2, we performed linear regression by using the model *y = ax*, where *x* represents the amplitude of each FBR component and *y* represents the amplitude of each component in the LR. As shown in Figure 6c, when FBR amplitudes were used to predict LR amplitudes across second shift locations (6, 11, 16 cm), the phasic component exhibited poor prediction power (left panel, *R*^2^ = −0.208), as did the tonic component (right panel, *R*^2^ = −0.040). In contrast, when LRs were regressed against the tonic FBR amplitude (Fig. 6d), both the phasic (left panel, *R*^2^ = 0.978) and tonic (right panel, *R*^2^ = 0.971) components of the LR showed strong linear relationships. These findings indicate that the phasic FBR component does not reliably account for the LR. In contrast, the tonic FBR component reflected the combined effects of the first and second cursor shifts, and its summed magnitude closely predicted the amplitude of the subsequent LR.

### The tonic component of the FBR predicts across-subject variability in the amplitude of the LR

In Experiments 1–3, we characterized the relationship between FBRs and LRs across different temporal patterns and magnitudes of visual error. However, LRs could, in principle, be generated directly from visual error without relying on FBRs. Indeed, previous studies have shown that measurable LRs can occur even when only endpoint error is provided, in the absence of within-trial FBRs [13,15, 26–27]. To examine whether individual differences in LR amplitude were more closely related to the tonic component of the FBR than to the nominal visual error, we compared FBRs and LRs under an identical temporal pattern of visual error. All participants in Experiments 1–3 (N = 33) experienced a cursor shift perturbation in which the cursor was laterally shifted 3 cm at the 1 cm location from the starting position (Fig. 7a). We therefore performed an additional analysis using this common temporal visual error pattern to examine whether across-subject variability in LR amplitude was predicted by the tonic component of the FBR.

**Figure 7:**
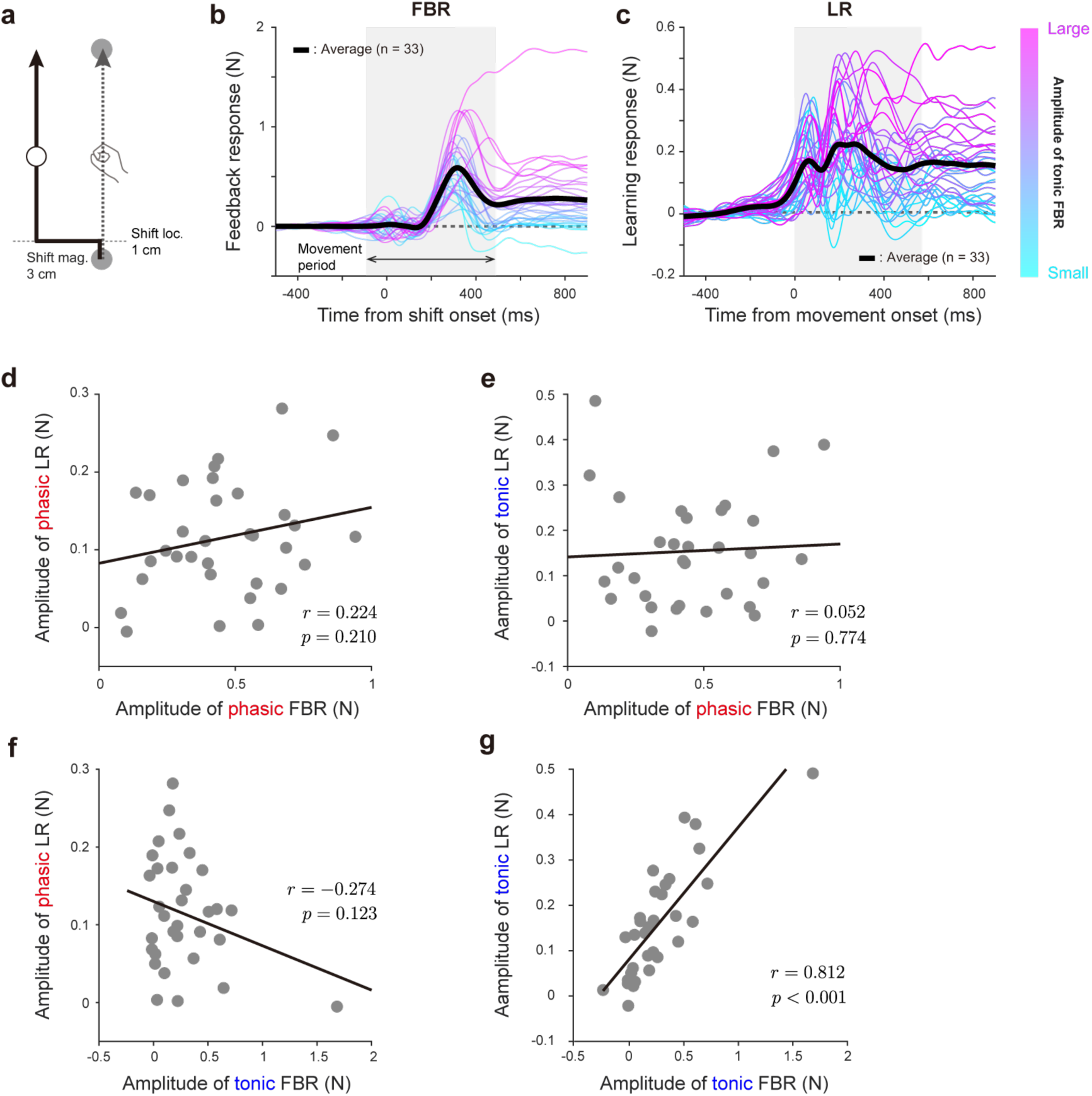
The tonic component of the feedback response predicts between-subject variability in learning response amplitude. **a**, Across Experiments 1–3, all participants experienced an identical temporal pattern of visual error, in which the cursor was shifted laterally by 3 cm at a location 1 cm from the starting position. **b-c**, Temporal profiles of the feedback response (FBR) (**b**) and learning response (LR) (**c**) elicited by this perturbation. Each trace represents an individual participant and the thick black trace represents the average across participants. Participants are ordered according to the amplitude of the tonic component of the FBR, with colors closer to magenta indicating larger tonic feedback responses. **d-g,** Relationships between the amplitudes of the phasic and tonic components of the FBRs and LRs across participants.

Figures 7b and 7c show the temporal profiles of the FBR (Fig. 7b) and LR (Fig. 7c) elicited by visual errors of the identical magnitude. Participants who exhibited larger FBRs during the holding period tended to produce larger LRs. To quantify these individual differences, we estimated the amplitudes of the FBRs and LRs by fitting their temporal profiles to the decomposition model (FBR: *R*^2^ = 0.9923, σ= 62, *T* = 300; LR: *R*^2^ = 0.9697, σ = 138, *T* = 136) and compared the extracted component amplitudes across participants. To directly assess whether individual differences in LR generation can be predicted from FBRs, we examined correlations between the amplitudes of the phasic and tonic components of the two responses (Figs. 7d-g). No significant correlation was observed between the phasic component of the FBR and either the phasic component of the LR (Fig. 7d; phasic, r = 0.224, p = 0.210) or tonic component of the LR (Fig. 7e; r = 0.052, p = 0.774). Similarly, the tonic component of the FBR did not correlate with the phasic component of the LR (Fig. 7f; r = -0.274, p = 0.123). In contrast, a strong and significant positive correlation was observed between the tonic components of the FBRs and LRs (Fig. 7g; r = 0.816, p < 0.001). Together, these results indicate that, at the level of individual differences, the tonic component of the FBR selectively predicts the amplitude of the tonic component of the LR.

### The tonic component of the FBR predicts trial-by-trial variability in LR amplitudes

We next examined trial-by-trial variability in LR amplitudes. Even when the same visual error was applied, some trials elicited larger LRs whereas others elicited smaller ones. We tested whether this variability could be predicted from the amplitude of the tonic component of the FBR using the data of the same condition in Experiments 1–3 (±3 cm cursor shift at the 1 cm location; N = 33). For each participant, trials were randomly divided into two datasets. These datasets were classified as High-FBR or Low-FBR based on the amplitude of the tonic component extracted from the averaged FBR. During this procedure, we ensured that the baseline trial force profiles were comparable between the two datasets, because differences in baseline force profiles can strongly influence the classification (see Methods). This random partitioning and selection procedure was repeated 100 times for each participant, and the resulting High-FBR and Low-FBR waveforms, as well as their corresponding LR waveforms, were obtained by averaging 100 waveforms. The decomposition model (Eq. 1) was then applied to the averaged waveforms to calculate the amplitudes of the phasic and tonic components of FBRs and LRs.

Figure 8a shows the force waveforms corresponding to the Low- and High-tonic FBR components, averaged across all participants in Experiments 1–3 (N = 33). Clear differences were observed during the late period between the two datasets, confirming successful partitioning. Figure 8b summarizes the difference in the amplitudes of the phasic and tonic components of the FBR between High- and Low-FBR datasets for all 33 participants. A one sample t-test confirmed that this classification produced a significant difference only in the tonic component (Fig. 8b; t(32) = 8.737, p < 0.001), with no significant difference in the phasic component (Fig. 8b; t(32) = 1.192, p = 0.242). The corresponding temporal profiles of the LR are shown in Figure 8c. A significant difference was observed in the amplitude of the tonic LR component between the two datasets (Fig. 8d; t(32) = 3.785, p < 0.001), corresponding to an average 16.76% reduction in tonic LR amplitude in the High-tonic FBR dataset relative to the Low-tonic FBR dataset. In contrast, no significant difference was observed in the phasic component of LR (Fig. 8d; t(32) = 0.278, p = 0.783). Importantly, these differences cannot be attributed to differences in baseline force waveforms, as the force waveforms of the baseline trials were comparable between the two datasets (Figs. 8a, c).

**Figure 8:**
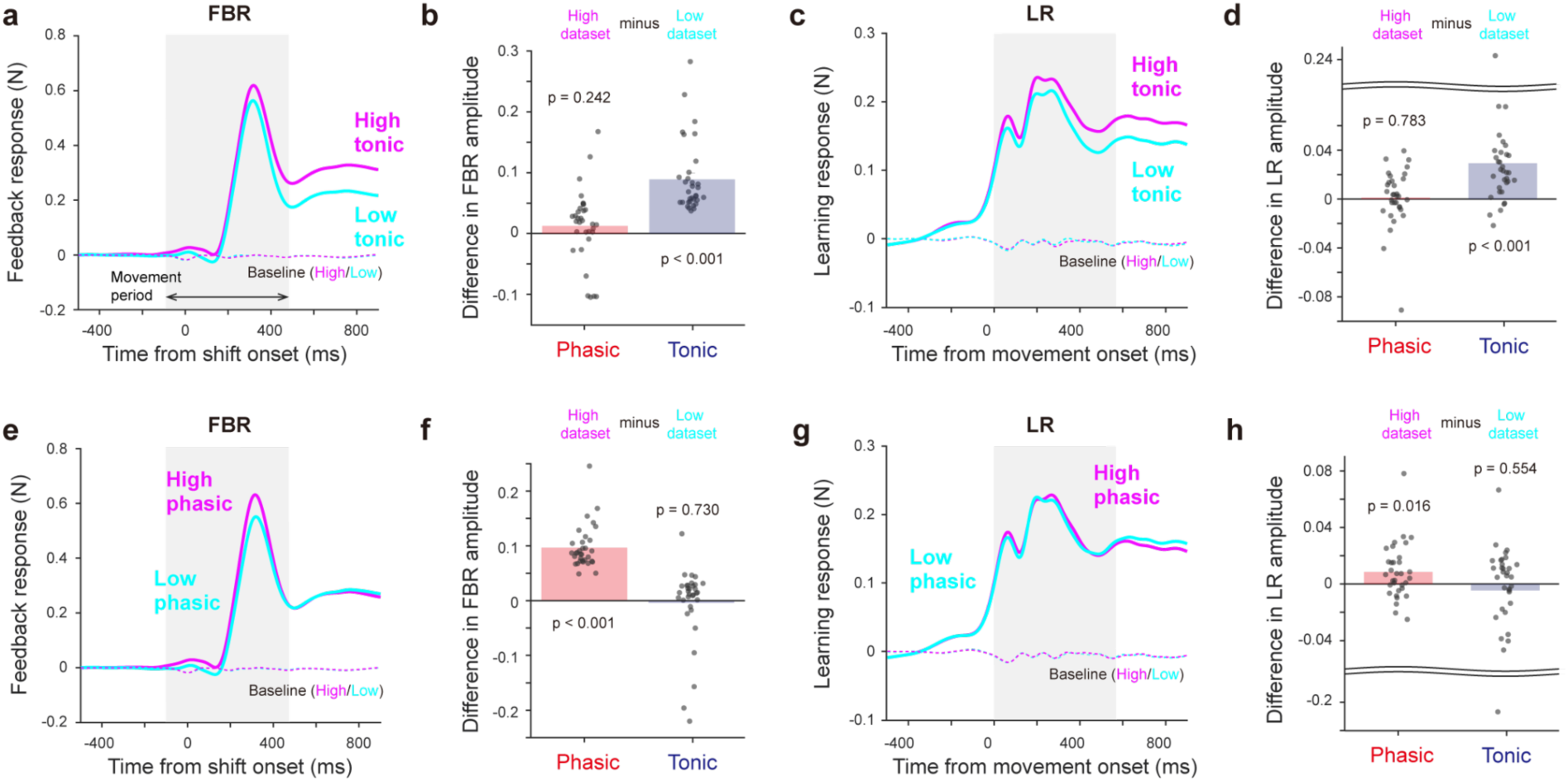
The tonic component of the feedback response predicts trial-by-trial variability in learning response amplitude. Trial triplets obtained under an identical cursor shift perturbation (see Fig. 7a) were split into two datasets based on the amplitude of either phasic (**a-d**) and tonic (**e-h**) components of the feedback response (FBR). **a**, Temporal profiles of the FBR for datasets split according to the tonic FBR component and averaged across participants. **b**, Difference in the amplitudes of the phasic and tonic components of the FBR between the High and Low tonic FBR datasets. **c-d**, Temporal profiles (**c**) and difference in the amplitudes of the phasic and tonic components (**d**) of the learning response (LR) for datasets split based on the tonic FBR component. **e,** Temporal profiles of the FBR for the datasets split according to the phasic FBR magnitude. **f,** Amplitudes of the phasic and tonic FBR components for the High- and Low phasic FBR datasets. **g–h,** Temporal profiles (**g**) and component amplitudes (**h**) of the LR for datasets split according to the phasic FBR amplitude.

We next repeated this analysis by classifying trials based on the amplitude of the phasic component of the FBR (Fig. 8e). As expected, this classification yielded significant differences only in the phasic component of the FBR (Fig. 8f; t(32) = 14.29, p < 0.001), with no differences in the tonic component (Fig. 8f; t(32) = -0.349, p = 0.730). However, classification based on the phasic FBR component produced little change in the temporal profile of the LR (Fig. 8g). No significant difference was found in the amplitude of the tonic LR component between the two datasets (Fig. 8d; t(32) = -0.60, p = 0.55, one-sample t-test). Although classification based on the phasic FBR component yielded a small but statistically significant difference in overall LR amplitude (t(32) = 2.54, p = 0.016), this effect was modest (6.15% change) and substantially smaller than effect of the tonic-FBR–based classification on the tonic LR component (16.76%). Together, these results indicate that the tonic component of the FBR provides a stronger predictor of trial-by-trial variability in LR amplitude than does the phasic component.

## Discussion

Accurate movement relies on the motor system’s ability to update motor commands across trials based on sensory feedback about movement errors. A central unresolved question is how the motor system utilizes the temporal structure of sensory error information to generate the LR. According to the widely discussed feedback-error learning hypothesis [9], the FBRs produced within a trial serve as teaching signals for generating LRs. Although this hypothesis has received substantial support, it remains debated whether LRs are primarily constructed from FBRs or instead directly from visual error information, and how these different sources contribute to distinct features of the LR, such as its amplitude and temporal pattern.

To address these issues, we employed an experimental paradigm originally developed to investigate FBRs [16]. This approach was well suited for our purpose, as it allowed direct comparison of FBRs and LRs across a wide range of visual error patterns by measuring the lateral forces generated against a clamp channel. Our experimental results demonstrate that the motor system does not transfer the temporal profile of the FBR to the LR. Instead, it adjusts the amplitude of the LR, whose temporal pattern remains relatively invariant, according to the amplitude of the FBR, particularly the component that persists through the holding phase (i.e., tonic component). These findings identify the tonic component of the FBR as a robust predictor of LR amplitude and motivate the hypothesis that sustained feedback-related signals contribute to subsequent motor-command updates. We return to this hypothesis below and discuss whether this relationship can be interpreted as causal in the sense proposed by the feedback error learning hypothesis.

### The LR is not a fixed time-shifted version of the FBR

A fundamental problem in motor control is how the motor system updates motor commands in response to movement errors. This updating process is nontrivial because the error information is represented in the sensory space and must be transformed into motor commands. The feedback error learning hypothesis offers an elegant solution to this problem by proposing that the FBR to a movement error—namely, the motor commands used to correct the error within a trial—serves as a teaching signal for generating the LR, expressed as updated motor commands in the subsequent trial.

A direct prediction of this hypothesis is that the LR should resemble a time-shifted version of the FBR. Supporting this view, Albert and Shadmehr [10] reported that, under a velocity-dependent force-field perturbation, the LR closely matched a fixed time-shifted copy of the FBR. They suggested that the magnitude of this time shift, approximately 125 ms, reflects the sensorimotor delay required for the motor system to generate the FBR in response to movement errors. However, because their study examined only a single type of force-field perturbation, it remains unclear whether the motor system truly shifts the temporal profile of the FBR forward in time based on sensory processing delays.

Addressing this question requires systematic manipulation of error timing. Fine and Thoroughman [11] pursued this approach by applying brief force pulses at different positions along the reaching trajectory. They found that FBRs emerged with a short latency after perturbation onset, whereas LRs in the subsequent trial consistently appeared near movement onset, largely independent of perturbation timing. However, the range of onset timings tested in that study was relatively narrow (approximately 200 ms), and both FBRs and LRs were inferred from movement trajectories, which may not sensitively reflect changes in underlying motor commands. These methodological constraints may have obscured differences in responses to force pulses applied at different times.

In the present study, we combined cursor-shift perturbations with an error-clamp method, allowing visual errors to be imposed precisely while minimizing changes in movement kinematics and avoiding secondary errors induced by FBRs or LRs themselves. This approach enabled direct comparison of the temporal profiles of the visual errors, FBRs, and LRs (Fig. 1). By systematically manipulating the onset of the cursor shift (Experiment 1), we found that the temporal relationship between FBRs and LRs was not fixed (Fig. 2f). Even when visual error onset varied widely—from shortly after movement onset to just before movement offset—the LR consistently emerged before the movement onset. Consistent with the findings of Fine and Thoroughman [11], these results demonstrate that the LR does not preserve temporal information about when the error occurred and therefore cannot be considered a fixed time-shifted copy of the FBR.

Notably, this lack of temporal specificity in the learning response contrasts with classical forms of cerebellar associative learning, such as delay eyeblink conditioning, in which the cerebellum acquires precise timing relationships between conditioned and unconditioned stimuli [28]. Moreover, recent work in mice has demonstrated temporally specific adaptation and aftereffects depending on the position of mossy fiber stimulation during reaching movements [29], where stimulation acts as an unconditioned signal. The temporal specificity observed in these paradigms differs markedly from the present findings in human visuomotor adaptation. Further studies will be required to clarify which cerebellar mechanisms distinguish temporally precise associative learning from the form of adaptation examined here.

### Nonlinear modulation of LR amplitude with error size parallels that of the tonic component of the FBR

Experiment 2 examined the quantitative relationship between error magnitude and both FBRs and LRs by systematically manipulating the magnitude of visual errors. Previous studies have shown that the LR amplitude increases approximately linearly for small errors but saturates for larger errors [30–33]. Several studies have also reported differential modulation of FBR and LR by error magnitude [11, 13, 34]. However, in most of these studies, the visual error experienced by participants could change during the movement as a consequence of the FBR itself, complicating precise characterization of the temporal evolution of the FBR.

The cursor-shift perturbation paradigm, which imposes a constant visual error throughout the movement [16], is therefore particularly well suited for examining how error magnitude modulates FBRs and LRs. Using this approach in Experiment 2, we found that FBRs were more sensitive to changes in error magnitude than LRs (Figs. 3b–c). Notably, consistent with findings using brief (approximately 250 ms) cursor-shift perturbations [23], this heightened sensitivity was confined to the early component of the FBR and was largely absent in the late component.

Motivated by previous work suggesting that motor commands can be decomposed into phasic and tonic components [25] or into velocity- and position-dependent components [24], we decomposed both FBRs and LRs into two analogous components and examined how each scaled with error magnitude (Figs. 4a–d). The phasic component of the FBR increased approximately linearly with error magnitude up to 2.0 cm (Fig. 4e), whereas the other three components—the tonic FBR and both components of the LR—exhibited similar nonlinear modulation, saturating even at small error magnitudes (Figs. 4e–h). Because our paradigm enabled full temporal profiling of both FBRs and LRs, we were able to identify these component specific scaling relationships directly.

Although numerous computational models have been proposed to account for the nonlinear modulation of LR amplitude with error magnitude [13, 30–33], our findings show that such nonlinearities already emerge during the later phase of the FBR. The close correspondence between the tonic FBR and LR is consistent with the possibility that these responses rely on partially overlapping computations [34, 35–36].

### LRs differ from FBRs in temporal pattern and vary primarily in amplitude

The temporal profiles of FBRs and LRs in Experiments 1 and 2 showed partial similarity, consistent with the findings of Albert and Shadmehr [10]. However, this similarity may reflect the use of simple step-like cursor shift perturbations, which impose limited temporal structure on the visual error. In contrast, Wei et al. [12] examined a broader range of force perturbations and evaluated FBRs and LRs using movement kinematics. They reported that FBRs exhibited temporally specific corrections that closely followed the time course of the applied force, whereas LRs displayed relatively nonspecific temporal profiles. However, because their analyses relied on movement trajectories, it was not possible to isolate the temporal evolution of the underlying motor commands or the sensory error from the corrective forces generated during movement.

As described above, our experimental paradigm avoids these confounds by imposing temporally precise visual errors while constraining movement kinematics, thereby enabling direct comparison of the motor commands underlying FBRs and LRs. When visual perturbations containing double cursor shifts were introduced in Experiment 3 (Fig. 5a), FBRs closely track the temporal structure of the imposed error. In contrast, LRs neither reflected the temporal pattern of the visual error nor resembled the temporal profile of the FBR (Figs. 5b–d). Instead, the motor system generated a temporally nonspecific LR that emerged immediately before movement onset (Figs. 5e–g). These results provide strong evidence that LRs do not inherit the temporal structure of FBRs, but instead differ primarily in amplitude. This finding extends and strengthens the conclusions of Wei et al. [12] by demonstrating this dissociation at the level of motor commands rather than inferred kinematics.

### The tonic component of the FBR as a candidate teaching-related signal for LR amplitude

The results of Experiment 2 revealed a close quantitative relationship between the tonic component of the FBR and the amplitude of the LR. This relationship was supported by three complementary analyses. First, inter-individual differences in LR magnitude were significantly correlated only with the amplitude of the tonic component of the FBR (Fig. 7), consistent with the previous findings of Albert and Shadmehr [10], who examined EMG activity associated with FBRs and LRs. Second, trial-by-trial variability in LR amplitude was explained by the tonic component of the FBR, but not by the phasic component (Fig. 8). Third, in Experiment 3, the amplitude of the LR induced by the second cursor shift was also predicted by the amplitude of the tonic component of the FBR evoked by the second cursor shift (Fig. 6).

An important issue is whether the close quantitative relationship between the tonic FBR and the LR can be interpreted as evidence for a causal influence of the FBR on the LR. This relationship alone does not necessarily establish that the FBR contributes directly to generating the LR, as proposed in the feedback error learning hypothesis. One alternative interpretation is that visual error information, combined with proprioceptive information, is used to estimate the hand deviation induced by the cursor perturbation, and that this estimated deviation contributes to both the FBR [37] and the LR [26, 38–39]. Under this interpretation, the relationship between the FBR and LR would reflect a shared state estimate rather than a causal influence of the FBR on the LR.

However, our trial-by-trial analysis revealed a relationship that cannot be fully explained by the nominal visual error alone. As shown in Figure 8, even when the imposed visual error was identical across trials, variability in the magnitude of the tonic FBR component was associated with variability in the subsequent LR. Moreover, this relationship was selective for the tonic component, whereas variability in the phasic FBR component had only a modest effect on the LR. Together, these findings support the interpretation that the tonic FBR is not merely a passive correlate of the imposed visual error, but may participate in the process that scales the subsequent LR.

### Origin of the tonic FBR component

Recent work suggests that the tonic component sustained until movement offset reflects an integrated motor command accumulated over the course of the movement [25]. Albert and colleagues [25] demonstrated an integrative linkage between moving and holding commands, showing that the time integral of movement-related force persists into the holding period and contributes to postural control. Similar integration mechanisms have been identified across multiple neural systems. In the oculomotor system, sustained neural activity following saccade termination has been attributed to a neural integrator that accumulates transient movement-related signals to maintain gaze [40–41]. In spatial navigation, neurons in the medial entorhinal cortex integrate velocity and directional inputs over time to estimate position [42–43]. Across systems, such integrative mechanisms transform temporally distributed signals, such as velocity, into representations of the current state [40]. In this framework, the feedback controller may integrate the feedback response itself, which reflects the temporal pattern of sensory error, to infer the state of the environment. Under this framework, the resulting tonic FBR may constitute a candidate teaching-related signal, in the sense proposed by the feedback error learning hypothesis, that is closely linked to the scaling of the LR on the subsequent trial.

### Perspectives for future studies

Despite the present findings, important aspects of how LRs are generated remain unresolved. First, the temporal structure of the visual error used in this study was relatively simple, consisting of linear combinations of stepwise cursor shifts. It remains to be determined whether the relationship between the tonic component of the FBR and the amplitude of the LR generalizes to more complex structures—such as nonlinear, nonstationary, or stochastic perturbations. Establishing the reproducibility of this mechanism across a broader class of error dynamics will be essential for evaluating its generality.

Second, the temporally nonspecific nature of the LR observed in the present study does not fully account for previous findings showing that the motor system can acquire complex, time-varying motor commands, such as those required to adapt to S-shaped force perturbations [25, 44]. How trial-by-trial updating of motor commands gives rise to such structured LRs under more dynamic conditions remains an important open question.

Finally, the present study does not address learning mechanisms that operate in the absence of online FBRs. It is well established that endpoint (terminal) visual feedback can induce LRs without engaging FBRs during the movement [13, 26–27], and that motor imagery can also drive learning in response to visual error [45]. Although endpoint error is generally considered a task error rather than a sensory prediction error and therefore engage reinforcement-like mechanisms distinct from error-based adaptation [46–47], an important challenge for future work will be to clarify how LRs are generated from these fundamentally different sources of error.

### Summary

In summary, we characterized the relationship between feedback responses and learning responses using an experimental paradigm that enabled direct comparison of their temporal profiles across multiple patterns of visual error. Our results show that the motor system does not transfer the temporal structure of the feedback response to the learning response. Instead, the amplitude of the tonic component of the feedback response—which reflects the temporal history of the visual error—strongly predicts the magnitude of the learning response. Moreover, the amplitude of this tonic component accounted for both inter-individual and trial-by-trial variability in learning response amplitude. These findings identify a robust quantitative link between sustained feedback-related responses and subsequent motor-command updates and provide evidence suggestive of a causal contribution of the tonic feedback response to learning.

## Methods

### General experimental procedures

A total of thirty-four right-handed participants (20 men and 14 women; age range, 18–40 years) volunteered to take part in the study. Each participant completed one of the three experiments. Participants were recruited using an online recruitment system (https://www.jikken-baito.com). All participants provided written informed consent prior to participation, and all experimental procedures were approved by the Office for Life Science Research Ethics and Safety of the University of Tokyo.

Participants performed planar reaching movements with their right hand while holding the handle of a robotic manipulandum (KINARM End-Point Lab, Kinarm, Kingston, Canada). A horizontal screen positioned above the handle displayed a starting position, a target (5 mm in diameter), and a white cursor (5 mm in diameter) indicating the handle position. This screen occluded direct vision of the participant’s arm and the handle.

The target was displayed 20 cm directly in front of the starting position. Approximately one second after the participant placed the handle at the starting position, the target changed color to magenta, serving as the “go” cue. Participants were instructed to move the handle smoothly and as straight as possible toward the target and to avoid using explicit strategies, even when the cursor was perturbed. The cursor disappeared 700 ms after movement completion (i.e., during the holding period), and the handle was automatically returned to the starting position. Participants were instructed to hold the handle at the perceived location of the target and to refrain from moving during the holding period.

To maintain consistent movement speed across trials, feedback on peak velocity was provided at the end of each trial: “too fast” (> 750 mm/s), “rather fast” (> 650 mm/s), “rather slow” (< 450 mm/s), or “slow” (< 350 mm/s). Before the main experiment, participants completed 50 practice trials to stabilize their movement accuracy and velocity.

### Experiment 1: Effect of visual perturbation onset on the LR

In Experiment 1 (10 participants: seven men and three women), we employed a single-trial visuomotor adaptation paradigm using a visual cursor-shift perturbation originally developed to investigate FBRs [16]. In perturbation trials, the cursor was shifted laterally by ±3 cm at one of four distances from the starting position (Fig. 2a; 1, 6, 11, and 16 cm). A baseline condition with no cursor shift was also included. Each perturbation trial was flanked by channel trials (Fig. 1a). During both perturbation and channel trials, hand trajectories were constrained to a straight path from the starting position to the target using a clamp-channel method [18], allowing measurement of the lateral force exerted against the channel. The force channel was implemented using a virtual spring (6000 N/m) and damper [100 N/(m/s)].

The temporal profile of the FBR was quantified as the difference in the lateral force between the perturbation and baseline trials, whereas the LR was quantified as the difference between the probe and baseline trials [10], During probe trials, the cursor was removed to assess the LR in the absence of visual feedback.

Each trial set consisted of five consecutive trials: a baseline trial to measure the baseline lateral force profile, a perturbation trial (one of nine conditions: 4 positions × 2 shift directions + 1 no shift condition), a probe trial (Fig. 1a), and two null trials without force channel or cursor perturbation to wash out adaptation effects. One cycle comprising all nine perturbation types was repeated 26 times. Including 50 practice trials, the total number of trials was 1,220 (50 practice trials + 5 trials × 9 perturbations × 26 cycles).

### Experiment 2: Effect of the visual error magnitude on the LR

Experiment 2 (12 participants: six men and six women) examined the quantitative relationship between FBRs and LRs across different magnitudes of cursor-shift perturbations. Participants repeatedly performed a set consisting of five trials. In perturbation trials, the cursor was shifted laterally by ±0.4, 0.8, 1.2, 1.6, 2.0, or 3.0 cm at a position 1 cm from the starting position (Fig. 3a). After 50 practice trials, one cycle comprising all 12 perturbation types was repeated 18 times, resulting in a total of 1,130 trials (50 practice trials + 5 trials × 12 perturbations × 18 cycles). One participant terminated the experiment early after completing 973 trials; for this participant, only the first 15 cycles were included in the analysis.

### Experiment 3: Effect of the complex temporal patterns of visual error on the FBR and LR

Experiment 3 (12 participants: seven men and five women) examined whether different temporal patterns of visual perturbations induce distinct patterns of FBRs and LRs. As in Experiments 1 and 2, participants repeatedly performed a set consisting of five trials. In the perturbation trials, three types of temporal perturbation patterns were used (Fig. 5a). In the maintained conditions, the cursor was shifted by ±3 cm at a location 1 cm from the starting position and the shift was maintained throughout the movement (2 types; +3cm or -3cm). In the removed conditions, the cursor was shifted by ±3 cm at the 1 cm location and then returned to zero (±3 cm → 0 cm) at one of three locations (6, 11, or 16 cm; 6 types). In the reversed conditions, the cursor was shifted by ±3 cm at the 1 cm location and then reversed (±3 cm → ∓3 cm) at one of the three locations (6, 11, or 16 cm; 6 types). In total, fourteen perturbation types were used. After 50 practice trials, one cycle comprising all 14 types was repeated 18 times, yielding 1,310 trials (50 practice trials + 5 trials × 14 perturbations × 18 cycles). One participant reported using an explicit strategy during the task and was therefore excluded from all analyses.

### Data analysis

Handle position and force data were sampled at 1000 Hz and low-pass filtered using a 4th-order zero-lag Butterworth filter with a cutoff frequency of 6 Hz. Handle velocity was calculated by numerically differentiating the position signal. The temporal profile of the FBR was defined as the difference between lateral force profiles measured during the perturbation and baseline trials, whereas the temporal profile of the LR was defined as the difference between lateral force profiles measured during the probe and baseline trials.

#### Cross-correlation analysis

Temporal similarity and relative timing between FBRs and LRs were evaluated using cross-correlation analysis. In Experiment 1, within-subject cross-correlations were computed to determine the time lag that maximized the correlation between the two responses. For Experiments 2 and 3, similarity was quantified using the cross-correlation coefficient at a fixed time lag corresponding to the peak of the averaged cross-correlation function obtained in Experiment 1 (cursor shift at 1 cm). In Experiment 2, cross-correlation coefficients were additionally computed at a longer fixed lag of -500 ms to assess differences in the overall shape of the cross-correlation functions.

#### Onset detection using ROC analysis

A receiver-operating characteristic (ROC) analysis was used to determine the onset latency of the FBR (Green & Swets, 1974; Nashed et al., 2014). ROC curves were constructed to quantify the probability of discriminating FBRs elicited by leftward versus rightward cursor shifts, with force profiles aligned to the onset of the cursor shift. The discrimination point was defined as the earliest time at which the ROC value exceeded 0.75 for at least 5 ms. To estimate the onset at which the ROC curve began to deviate from chance (0.5), we regressed the ROC values within the 15 ms window preceding the discrimination point and calculated the time at which this regression line intersected the baseline of the ROC curve.

#### Decomposition analysis

To quantitatively characterize FBRs and LRs, we fitted their temporal profiles using Eq. 1. FBRs were aligned to the cursor-shift onset (Fig. 3a), whereas LRs were aligned to movement onset (Fig. 3b). Model fitting was first performed on response profiles averaged across participants to identify the parameters σ and *T*. Using these parameters, individual response profiles were then refitted to estimate the amplitudes of the phasic and tonic components. The same procedure was applied to FBRs and LRs elicited by the second cursor shift in Experiment 3. Specifically, the decomposition model was fitted to differential feedback response waveforms, computed as the difference between the removed or reversed conditions and the maintained condition, and aligned to the onset of the second shift (Fig. 6a, inset). An analogous analysis was performed for differential learning response waveforms aligned to movement onset (Fig. 6b, inset).

#### Linear regression relating FBR and LR amplitudes

To quantify the relationship between the phasic and tonic components of the FBR and LR, we performed linear regression using the model *y = ax*, where *x* represents the amplitude of each FBR component and *y* represents the amplitude of the corresponding LR component. Model performance was evaluated using the coefficient of determination (*R²*) calculated between predicted and observed LR amplitudes. *R²* values were computed across all perturbation conditions to evaluate how well the phasic and tonic components of the FBR accounted for modulation of the corresponding components of the LR.

#### Dividing trials into Low and High tonic or phasic FBR components

Even when identical cursor shift magnitudes were applied, both FBRs and LRs exhibited substantial trial-by-trial variability. We therefore examined whether variability in the FBR was systematically related to variability in the LR. To this end, we analyzed trials from the same condition in Experiments 1-3 (±3 cm cursor shift at 1 cm location; N = 33) and divided them into two datasets based on whether they exhibited larger tonic or phasic FBR components. For each participant, trials with rightward and leftward cursor shifts were randomly split into two datasets, and baseline force profiles, FBRs, and LRs were obtained for each dataset by collapsing force waveforms across cursor shift directions.

A critical issue in this classification procedure was that differences in baseline force profiles could strongly influence both FBRs and LRs. Because FBRs and LRs were computed by subtracting baseline force waveforms from the perturbed and probe trial waveforms, respectively, large differences in baseline force between datasets could lead to classification driven by baseline force variability rather than by genuine differences in FBRs. In such cases, both the FBRs and LRs waveforms would be contaminated by baseline force differences. To mitigate this problem, we repeated the random trial partitioning procedure 100 times and selected the pair of datasets that exhibited the highest similarity between baseline force profiles, defined as the smallest mean squared difference between baseline force profiles. This approach ensured that subsequent comparisons primarily reflected variability in feedback responses rather than differences in baseline force (see Figs. 8a, 8c, 8e and 8g).

We then fitted the FBR waveforms averaged within each dataset using Eq. 1 to estimate the amplitude of the phasic and tonic components. Based on the amplitude of either the phasic or tonic component of the FBR, each dataset was classified as a Low-phasic (or tonic) FRB or High-phasic (or tonic) FBR dataset. This procedure was repeated 100 times, and the baseline force, FBR and LR waveforms were averaged across repetitions for each Low and High FBR datasets. Finally, the grand averaged waveforms were again fitted using Eq. 1 to estimate the phasic and tonic components of both FBRs and LRs.

#### Data exclusion

We excluded the data from the analysis by the following criteria. The lateral force data in each cycle was discarded when the peak movement velocity for either a baseline trial, a perturbation trial, or a probe trial was outside the included range (< 450 mm/s or >750 mm/s). The proportion of the excluded cycle was 2.52% (Experiment 1), 2.70 % (Experiment 2), and 3.25 % (Experiment 3).

## Acknowledgments

We thank members of the Nozaki Laboratory for their helpful comments and suggestions, as well as for their assistance in coordinating the experiments. This study was supported by grants from the Japan Society for the Promotion of Science (JSPS) Research Fellowships for Young Scientists (22KJ0992) to YM, JSPS KAKENHI (20J13734, 23K10739) to TK and JSPS KAKENHI (21H04860, 22K19736, 23K27958) to DN.

## Author contributions

Conceptualization, Methodology, Formal analysis, Investigation, and Writing: Y.M., T.K., D.N.; Experimental setup: Y.M.; Supervision: D.N.

## Data availability

The source Matlab data that support the findings of this study and the programming codes to analyze the data are available in Zenodo (10.5281/zenodo.18297833), and raw experimental data is available from the corresponding author upon reasonable request.

## Supplement Information

**Supplemental Figure 1:**
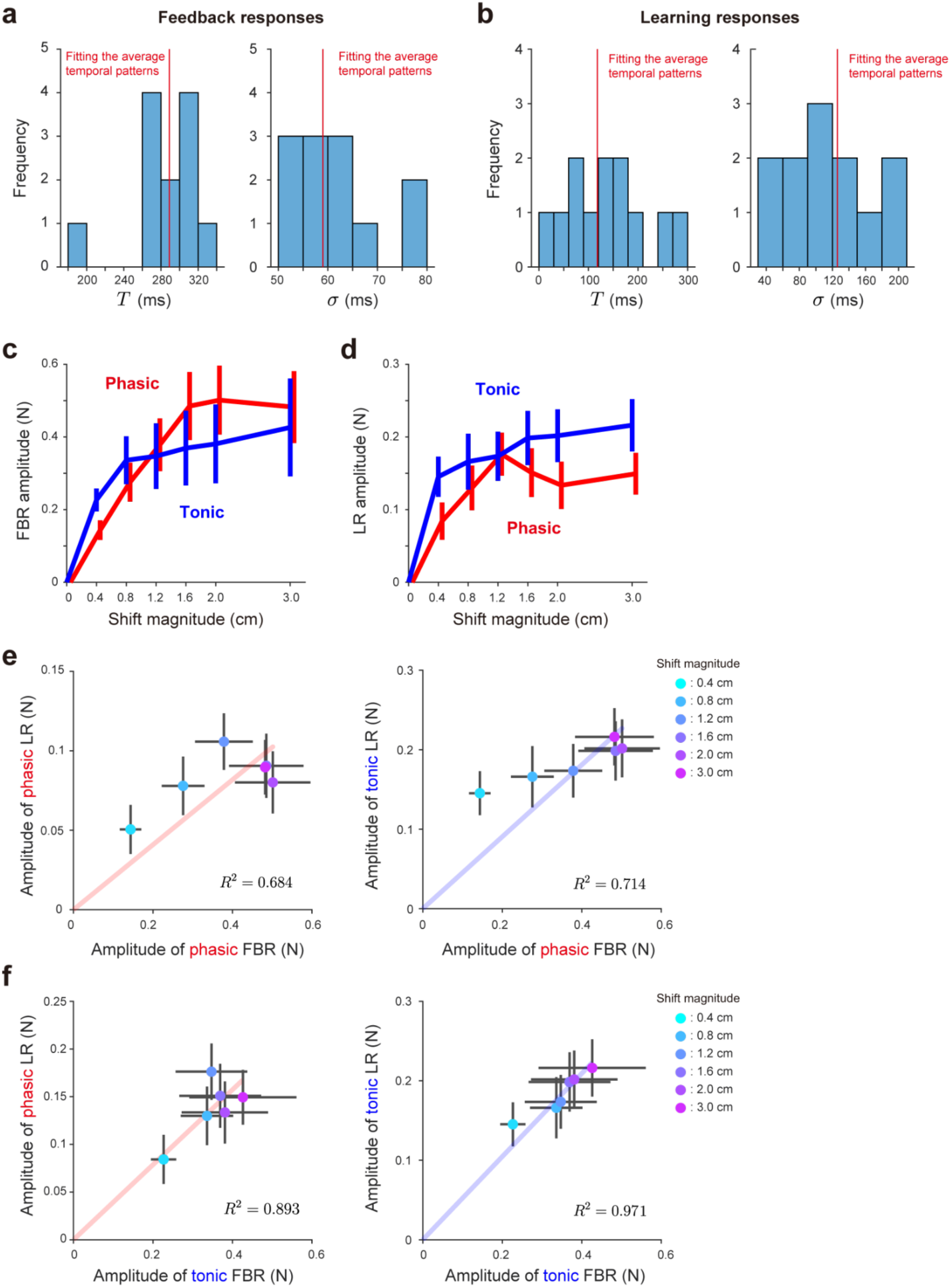
Robustness of the quantitative relationship between feedback and learning responses to model parameterization in Experiment 2. Even when additional fitting parameters were introduced for the model (Eq. 1), the quantitative relationship between the feedback response (FBR) and the learning response (LR) remained unchanged. **a-b,** The decomposition model (Eq. 1) was fitted to individual participants’ FBRs (**a**) and LRs (**b**) while allowing not only component amplitudes but also onset timing and temporal dispersion to vary as free parameters. The fitted parameter values across participants were distributed around those obtained from the grand-averaged responses. This distribution was relatively narrow for the FBR, indicating high consistency across participants, whereas the LR showed a broader distribution, reflecting greater inter-individual variability. **c-f,** Using this extended fitting procedure, the model successfully reproduced the modulation of FBR and LR amplitudes as a function of visual error magnitude, consistent with the results shown in Fig.4.

## References

1. Taylor JA, Ivry RB. Flexible cognitive strategies during motor learning. PLoS Comput Biol. 2011;7(3):e1001096.

2. Herzfeld DJ, Vaswani PA, Marko MK, Shadmehr R. A memory of errors in sensorimotor learning. Science. 2014;345:1349–1353.

3. Thoroughman KA, Shadmehr R. Learning of action through adaptive combination of motor primitives. Nature. 2000;407:742–747.

4. Donchin O, Francis JT, Shadmehr R. Quantifying generalization from trial-by-trial behavior of adaptive systems that learn with basis functions: theory and experiments in human motor control. J Neurosci. 2003;23:9032–9045.

5. Nozaki D, Kurtzer I, Scott SH. Limited transfer of learning between unimanual and bimanual skills within the same limb. Nat Neurosci. 2006;9:1364–1366.

6. Smith MA, Ghazizadeh A, Shadmehr R. Interacting adaptive processes with different timescales underlie short-term motor learning. PLoS Biol. 2006;4(6):e179.

7. Jordan MI, Rumelhart DE. Forward models: supervised learning with a distal teacher. Cogn Sci. 1992;16:307–354.

8. Jordan MI. Learning inverse mappings using forward models. In: Proceedings of the Sixth Yale Workshop on Adaptive and Learning Systems; 1990. p. 146–151.

9. Kawato M, Furukawa K, Suzuki R. A hierarchical neural-network model for control and learning of voluntary movement. Biol Cybern. 1987;57:169–185.

10. Albert ST, Shadmehr R. The neural feedback response to error as a teaching signal for the motor learning system. J Neurosci. 2016;36(17):4832–4845.

11. Fine MS, Thoroughman KA. Motor adaptation to single force pulses: sensitive to direction but insensitive to within-movement pulse placement and magnitude. J Neurophysiol. 2006;96(2):710–720.

12. Wei K, Wert D, Körding K. The nervous system uses nonspecific motor learning in response to random perturbations of varying nature. J Neurophysiol. 2010;104(6):3053–3063.

13. Makino Y, Hayashi T, Nozaki D. Divisively normalized neuronal processing of uncertain visual feedback for visuomotor learning. Commun Biol. 2023;6:1396.

14. Castro LNG, Hadjiosif AM, Hemphill MA, Smith MA. Environmental consistency determines the rate of motor adaptation. Curr Biol. 2014;24:1050–1061.

15. Hayashi T, Kato Y, Nozaki D. Divisively normalized integration of multisensory error information develops motor memories specific to vision and proprioception. J Neurosci. 2020;40(7):1560–1570.

16. Franklin DW, Wolpert DM. Specificity of reflex adaptation for task-relevant variability. J Neurosci. 2008;28:14165–14175.

17. Dimitriou M, Wolpert DM, Franklin DW. The temporal evolution of feedback gains rapidly update to task demands. J Neurosci. 2013;33:10898–10909.

18. Scheidt RA, Reinkensmeyer DJ, Conditt MA, Rymer WZ, Mussa-Ivaldi FA. Persistence of motor adaptation during constrained, multi-joint, arm movements. J Neurophysiol. 2000;84:853–862.

19. Green DM, Swets JA. Signal detection theory and psychophysics. Oxford: Robert E. Krieger; 1974.

20. Nashed JY, Crevecoeur F, Scott SH. Rapid online selection between multiple motor plans. J Neurosci. 2014;34(5):1769–1780.

21. Scott SH. A functional taxonomy of bottom-up sensory feedback processing for motor actions. Trends Neurosci. 2016;39(8):512–526.

22. Day BL, Lyon IN. Voluntary modification of automatic arm movements evoked by motion of a visual target. Exp Brain Res. 2000;130(2):159–168.

23. Franklin DW, Reichenbach A, Franklin S, Diedrichsen J. Temporal evolution of spatial computations for visuomotor control. J Neurosci. 2016;36:2329–2341.

24. Sing GC, Joiner WM, Nanayakkara T, Brayanov JB, Smith MA. Primitives for motor adaptation reflect correlated neural tuning to position and velocity. Neuron. 2009;64(4):575–589.

25. Albert ST, Hadjiosif AM, Jang J, Zimnik AJ, Soteropoulos DS, Baker SN, et al. Postural control of arm and fingers through integration of movement commands. eLife. 2020;9:e52507.

26. Wei K, Körding K. Uncertainty of feedback and state estimation determines the speed of motor adaptation. Front Comput Neurosci. 2010;4:11.

27. Burge J, Ernst MO, Banks MS. The statistical determinants of adaptation rate in human reaching. J Vis. 2008;8(4):20. doi: 10.1167/8.4.20.

28. Lavond DG, Knowlton BJ, Steinmetz JE, Thompson RF. Classical conditioning of the rabbit eyelid response with a mossy-fiber stimulation CS: II. Lateral reticular nucleus stimulation. Behav Neurosci. 1987;101:676–682.

29. Calame DJ, Becker MI, Person AL. Cerebellar associative learning underlies skilled reach adaptation. Nat Neurosci. 2023;26:1068–1079.

30. Wei K, Körding K. Relevance of error: What drives motor adaptation? J. Neurophysiol. 2009:101, 655–664.

31. Marko MK, Haith AM, Harran MD, Shadmehr R. Sensitivity to prediction error in reach adaptation. J Neurophysiol. 2012;108:1752–1763.

32. Kim HE, Morehead JR, Parvin DE, Moazzezi R, Ivry RB. Invariant errors reveal limitations in motor correction rather than constraints on error sensitivity. Commun Biol. 2018;1:19.

33. Tsay JS, Avraham G, Kim HE, Parvin DE, Wang Z, Ivry RB. The effect of visual uncertainty on implicit motor adaptation. J Neurophysiol. 2021;125:12–22.

34. Hayashi T, Yokoi A, Hirashima M, Nozaki D. Visuomotor map determines how visually guided reaching movements are corrected within and across trials. eNeuro. 2016;3(3):ENEURO.0032-16.2016.

35. Maeda RS, Cluff T, Gribble PL, Pruszynski JA. Feedforward and feedback control share an internal model of the arm’s dynamics. J Neurosci. 2018;38(49):10505–10514.

36. Maeda RS, Gribble PL, Pruszynski JA. Learning new feedforward motor commands based on feedback responses. Curr Biol. 2020;30(10):1941–1948.e3.

37. Körding KP, Wolpert DM. Bayesian integration in sensorimotor learning. Nature. 2004;427(6971):244–247.

38. Zhang Z, Wang H, Zhang T, Nie Z, Wei K. Perceptual error based on Bayesian cue combination drives implicit motor adaptation. eLife. 2024;13:e94608.

39. Tsay JS, Kim H, Haith AM, Ivry RB. Understanding implicit sensorimotor adaptation as a process of proprioceptive re-alignment. eLife. 2022;11:e76639.

40. Shadmehr R. Distinct neural circuits for control of movement vs. holding still. J Neurophysiol. 2017;117(4):1431–1460.

41. Robinson DA. Models of the saccadic eye movement control system. Kybernetik. 1973;14:71–83.

42. McNaughton BL, Battaglia FP, Jensen O, Moser EI, Moser MB. Path integration and the neural basis of the “cognitive map.” Nat Rev Neurosci. 2006;7:663–678.

43. Burak Y, Fiete IR. Accurate path integration in continuous attractor network models of grid cells. PLoS Comput Biol. 2009;5(2):e1000291.

44. Makino Y, Suemitsu K, Hirashima M. Action progress organizes motor memory. bioRxiv [Preprint]. 2026 [posted 2026 Feb 9; cited 2026 Mar 19]. Available from: https://www.biorxiv.org/content/10.64898/2026.02.09.704807v1 doi: 10.64898/2026.02.09.704807

45. Kim OA, Forrence AD, McDougle SD. Motor learning without movement. Proc Natl Acad Sci U S A. 2022;119(30):e2204379119.

46. Ikegami T, Ganesh G, Gibo TL, Yoshioka T, Osu R, Kawato M. Hierarchical motor adaptations negotiate failures during force field learning. PLoS Comput Biol. 2021;17(4):e1008481.

47. Kim HE, Parvin DE, Ivry RB. The influence of task outcome on implicit motor learning. eLife. 2019;8:e39882.

